# Reduced Dependence on Sensorimotor Processing in the Brain is Associated with Higher Math Skills in Adults

**DOI:** 10.1101/2025.09.16.676580

**Authors:** Xueying Ren, Marc N. Coutanche, Julie A. Fiez, Melissa E. Libertus

**Affiliations:** Department of Psychology, University of Pittsburgh, Pittsburgh, PA, USA 15260; Learning Research and Development Center, University of Pittsburgh, Pittsburgh, PA, USA 15260; Center for the Neural Basis of Cognition, Pittsburgh, PA, USA 15260; Brain Institute, University of Pittsburgh, Pittsburgh, PA, USA 15260

**Author notes:** Corresponding author: Name: Xueying Ren Tel: Address: 3420 Forbes Avenue, Murdoch, Pittsburgh, PA 15260, USA. The authors declare no competing financial interests.

**Keywords:** Numerical cognition, embodied cognition, disembodied cognition, math, fMRI

## Abstract

Number processing is crucial for math competence, yet the neural underpinnings remain unclear. In this study, we used functional magnetic resonance imaging (fMRI) to 1) identify key brain regions involved in symbolic and embodied number processing, and 2) examine their relations with math abilities in children and adults. We collected fMRI data from 104 adults (*M*_age_ = 23.25 years) and 88 4^th^ graders (*M*_age_ = 9.75) as they performed number comparison and phonological tasks with numeral (symbolic) and hand (embodied) numerical stimuli.

Participants’ math abilities were assessed using the math subtests of the Woodcock-Johnson Tests of Achievement. Univariate analyses revealed number processing regions in occipital, parietal and temporal cortices including inferior parietal, sensorimotor, supramarginal and insular areas. In adults, less involvement of sensorimotor processing regions during number compared to phonological processing is associated with enhanced math skills. These findings not only elucidate the neural mechanisms underlying number processing and math abilities but also raise questions about the emphasis on embodied number presentation in math learning experiences.

**Significance Statement:** Children and adults show substantial individual differences in math competence, and understanding the underlying neural mechanisms is crucial. By employing functional magnetic resonance imaging (fMRI), we identified key brain regions involved in symbolic and embodied number processing and explored their relations with math skills. Our findings reveal that reduced sensorimotor involvement during number processing correlates with enhanced math performance in adults. This study not only elucidates the neural mechanisms underlying number processing, offering critical insights into the cognitive architecture that supports math abilities, but also raises questions about the emphasis on embodied number representations in math learning experiences.

## 1. Introduction

### 1.1 The multi-faceted nature of number representations

Number processing underpins math competence (Menon, 2014). Symbolic integration, which involves linking numerical symbols with the quantities they represent, is a critical aspect of this process (Ansari et al., 2005; Lyons et al., 2012). According to Dehaene’s triple-code model (Dehaene, 1992), number processing incorporates three distinct but interacting codes: a verbal code for number words, a visual code for written digits and words, and a semantic code that conveys the meaning of numbers. Recent studies have proposed a potential fourth code—an embodied code, where especially fingers (but also other parts of the body) can represent quantities and influence number processing (Domahs et al., 2010).

Understanding the interrelations among numerical codes is crucial for cognitive development theory and educational applications. For instance, children’s ability to match and integrate non-symbolic and symbolic numerical information correlates with their school math performance (Mundy & Gilmore, 2009). Similarly, adults’ ability to integrate verbal and visual numerical information is associated with their arithmetic achievement (Sasanguie et al., 2017). However, recent behavioral evidence has suggested that the opposite may also be true. For example, greater symbolic estrangement (i.e., *less* integration) between visual and semantic codes is linked to better math skills in adults (Ren et al., 2022), and is unrelated to math abilities in children (Ren et al., 2025). Thus, the exact role of integrating different number codes is still unclear.

Different number codes are represented in different brain regions. The verbal code has been linked with the angular gyrus (Dehaene et al., 2003), the visual code with the fusiform gyrus (Abboud et al., 2015), the semantic code with the intraparietal sulcus (IPS; Hubbard et al., 2005), and the embodied code with sensorimotor cortices (Berteletti & Booth, 2015). The IPS is considered a hub for numerical integration, especially for quantities and symbolic numbers (Bugden et al., 2019; Fias et al., 2003; Koch et al., 2023). Infants process quantities in the right parietal area by six months (Hyde et al., 2010), and by age four, children engage the IPS for non-symbolic number processing, laying the foundation for symbolic understanding (Cantlon et al., 2006). The right IPS tunes to numerosity in early childhood (Edwards et al., 2016; Kersey & Cantlon, 2017), while the left IPS increasingly handles symbolic number processing and is linked to enhanced numerical abilities as children develop math skills (Ansari, 2016; Emerson & Cantlon, 2015).

### 1.2 The embodied number code

Numbers are traditionally viewed as abstract symbols, seemingly disconnected from any sensory or motor experiences (Piaget, 2013). However, theories of embodied cognition suggest that abstract number representations may be rooted in bodily experiences (Fischer & Shaki, 2018). Evidence shows that sensorimotor experiences during the acquisition of numbers play an important role in shaping numerical knowledge. For instance, a study found that embodied training, where first graders walk to estimated positions on a number line on the floor, enhances their understanding of numbers (Link et al., 2013). Also, finger differentiation training can improve children’s numerical abilities (Gracia-Bafalluy & Noël, 2008). Research into finger gnosis, i.e., the ability to differentiate individual fingers, has shown a clear association between finger gnosis and numerical skills in first graders (Noël, 2005). Moreover, early finger-related neuropsychological skills predict later math achievements. For example, Fayol et al. (1998) demonstrated that finger-related abilities at age five predict arithmetic competence at age six, suggesting that the ability to represent and discriminate fingers plays a key role in the development of quantity representation and problem-solving skills. Longitudinal studies reveal that finger counting is a critical, adaptive strategy in early numerical development. It is most beneficial when children are first learning number combinations, and its use naturally declines as their mental arithmetic skills increase (Jordan et al., 2008). In fact, proficiency in finger counting actively supports children’s math development such that kindergartners who use efficient finger strategies are more likely to transition successfully to mental calculation in later grades (Poletti et al., 2022). Childhood finger-counting habits continue to influence the mental representation and processing of numbers into adulthood (Di Luca et al., 2006; Di Luca & Pesenti, 2008). Finally, finger-based numerical representations can remain unconsciously active, influencing simple arithmetic operations even in adulthood (Badets et al., 2010).

Engaging the motor system through graphical representations or observed actions facilitates the understanding of abstract mathematical concepts, like numbers, by making them more tangible (Khatin-Zadeh et al., 2022). Converging neural evidence confirms this link, showing that numerical cognition is deeply rooted in the brain’s motor system. For instance, studies using positron emission tomography (PET) reveal an increase in the amplitude of motor-evoked potentials during counting tasks, indicating a close relation between hand and numerical representations (Sato et al., 2007). Other methods provide causal evidence; transcranial magnetic stimulation (TMS) has demonstrated the direct involvement of hand motor circuits during counting (Andres et al., 2007). Functional magnetic resonance imaging (fMRI) has further pinpointed the specific neural substrates of this connection. In children, counting processes activate the anterior intraparietal sulcus (aIPS), a region also engaged during visually guided finger movements (Krinzinger et al., 2011). This research has become even more specific, showing that different arithmetic operations rely on these circuits to varying degrees. For instance, subtraction problems, which require quantity manipulation, significantly activate finger motor and somatosensory areas in children, whereas multiplication problems do not. Interestingly, better subtraction performance was associated with lower activation in the somatosensory finger area, suggesting greater neural efficiency (Berteletti & Booth, 2015). In adults, the brain circuits involved in finger representation—including the superior parietal lobule and intraparietal sulcus—also underlie arithmetic operations (Andres et al., 2012). This enduring connection shows that even when processing abstract symbols, adults activate the specific perceptual-motor brain networks corresponding to their individual finger counting experiences, suggesting that number processing is deeply embodied (Tschentscher et al., 2012).

### 1.3 Current study

In this study, we employed a novel number localizer task to investigate the neural underpinnings of number processing in fourth-grade children and adults. We incorporated both symbolic and embodied codes to shed light on their roles for formal math competence at different development stages. We hypothesized that, compared to children, adults would leverage a broader set of brain regions for faster number processing. Furthermore, because adults have more experience with number processing, we hypothesized that adults with higher math skills would rely less on the embodied code, as indicated by less involvement of motor areas. Finally, as previous research has seldom addressed the specificity of correlations between brain activation and math ability, we tested whether number processing regions in the brain also correlate with reading skills.

## 2. Methods

### 2.1 Participants

A total of 106 adults and 97 children took part in this study, and written informed consent was obtained from all adult participants or legal guardians of child participants prior to any data collection. Adult data were collected from October 2018 to January 2020, and child data were collected from May 2018 to October 2023. All experimental procedures were approved by the local Institutional Review Board. All participants were native monolingual English speakers, right-handed, had normal or corrected-to-normal vision, and no history of learning disabilities. All participants were compensated monetarily for their participation; children also received a small gift (e.g., a book). For adults, two were excluded due to cysts discovered during the MRI scan. As a result, 104 adults (59 females, 45 males; age range = 18-35 years; mean age = 23.25 years, standard deviation (*SD*) = 4.5 years) remained in the final analyses. For children, one participant was excluded due to a cyst, and eight participants were excluded due to excessive overall head motion (> 5mm), resulting in 88 children (female = 38, male = 49, other = 1; age range: 8.17-10.83 years, mean age = 9.75 years, *SD* = 0.5 years) included in the final analysis.

### 2.2 fMRI tasks and stimuli

#### 2.2.1 Stimuli

In the scanner, we implemented a number localizer task with two types of stimuli: single-digit Arabic numerals (range 1-4 and 6-9), and hand images displaying numbers in iconic ways (e.g., an extended index finger for “1”). Stimuli were presented on a grey background at the center of the screen, with Arabic numerals in white color, and hand images as color photographs of actual human hands. For small numbers (< 5), the hand on the left side of the image was closed in a fist to indicate 0; for large numbers (> 5), the hand on the right side of the image was open to indicate 5 (see Fig. 1 for illustration).

**Fig. 1:**
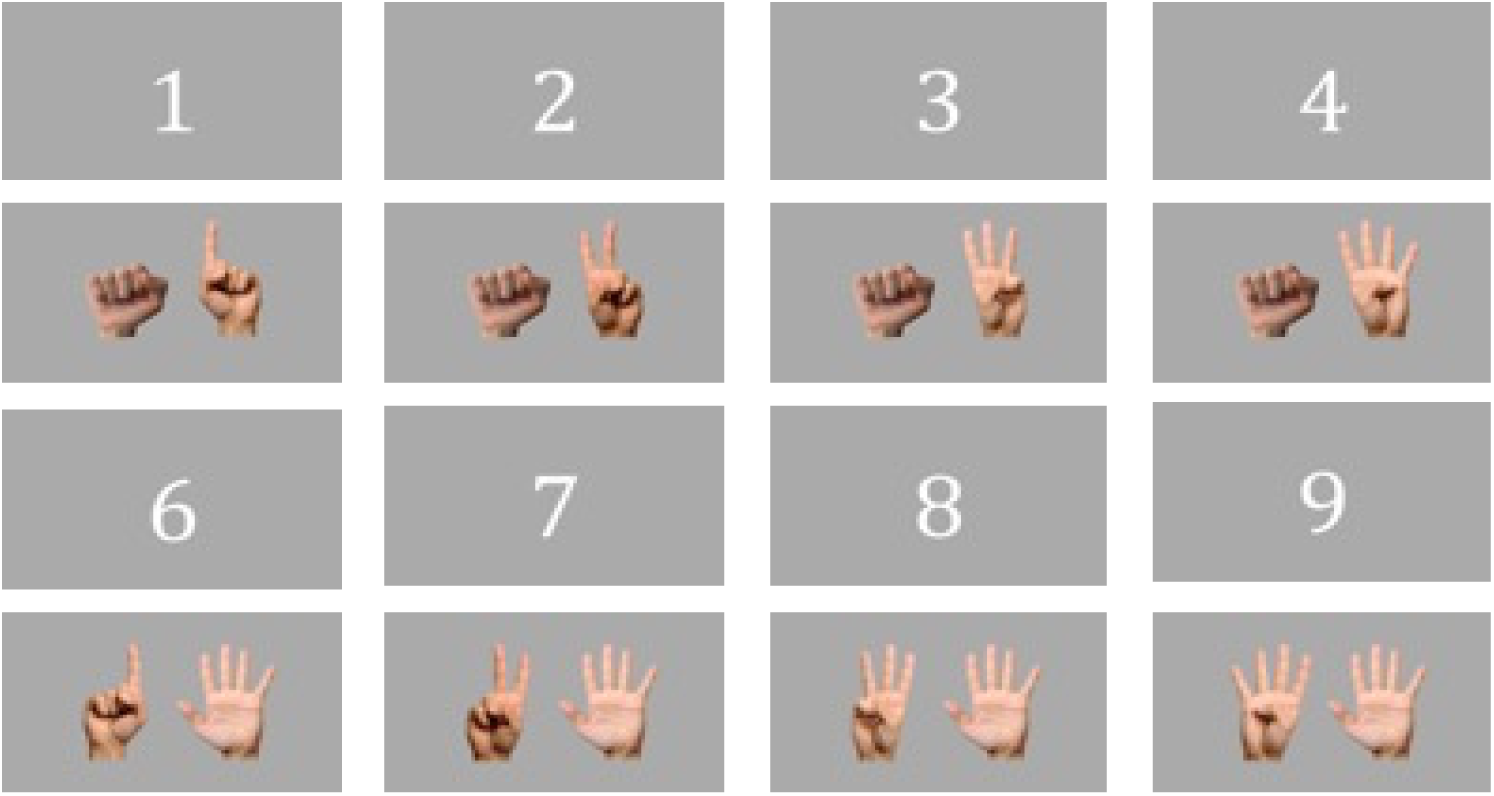
Two stimulus types used during the fMRI number localizer task: single-digit Arabic numerals and hand images. For small numbers (<5), the hand on the left side of the image was closed in a fist to indicate 0. For large numbers (>5), the hand on the right side of the image was open to indicate 5. Hand images included the iconic representations shown here (e.g., ‘8’ indicated by three and five (but not two sets of four) fingers raised).

#### 2.2.2 Task design

Participants were asked to complete two different tasks: a number and a phonological task. In the *number task*, participants were required to determine whether a number stimulus was larger or smaller than a target number (3, 4, 6, or 7) regardless of whether numerical information was presented as an Arabic numeral or a hand display. Target numbers were counterbalanced across blocks. In the *phonological task*, participants were asked to judge whether the first sound of the corresponding number word was the same as the first sound of the corresponding word of a target cartoon object (fan, nose, sun, tape; see Fig. 2 for illustration). Following the display of the stimulus, participants were asked to respond as accurately and quickly as possible before the end of a subsequent fixation cross.

**Fig. 2:**
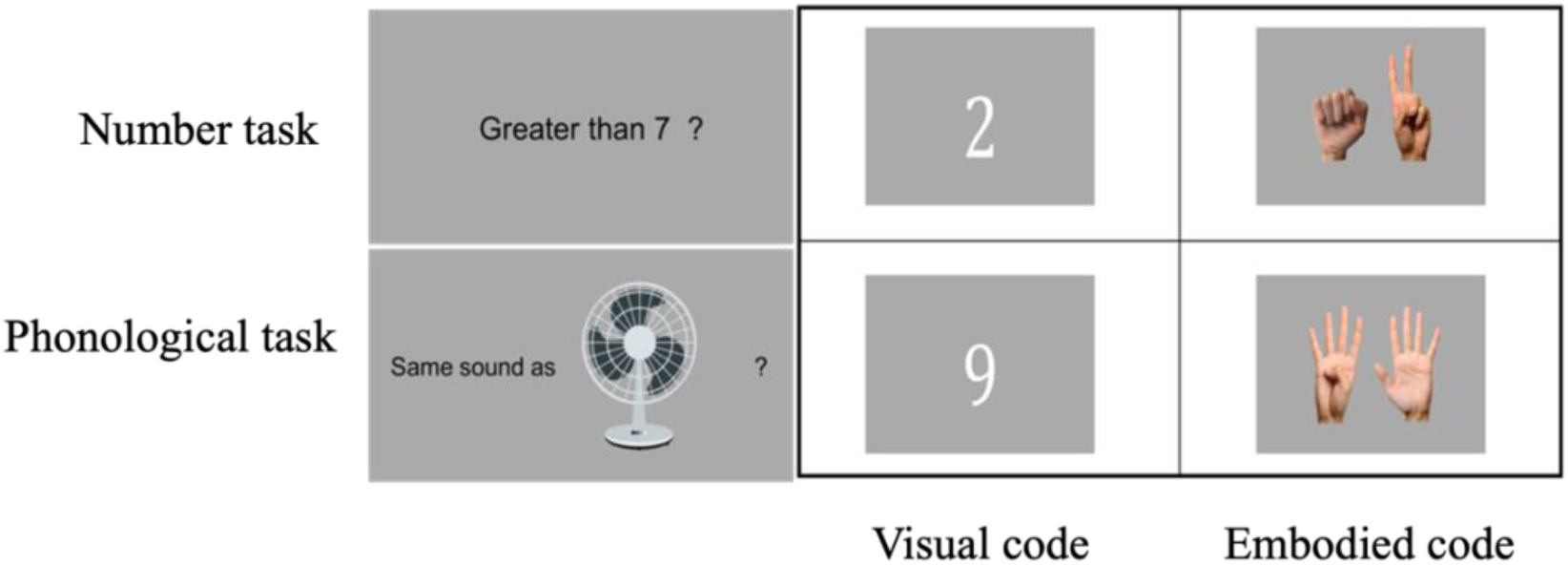
fMRI task design illustration. Two types of stimuli, Arabic numerals (visual code) and hand images (embodied code), were crossed with two types of tasks, number and phonological tasks, to generate four conditions.

We used a block design with a total of six runs. Each run consisted of eight blocks, i.e. two blocks for each of the four conditions. In each block, a prompt question was presented first lasting for eight seconds, followed by three trials of the same condition (e.g., a phonological task with Arabic numerals). Each trial lasted for two seconds, followed by two seconds of a fixation cross, resulting in 20 seconds per block (see Fig. 3 for illustration). The order of the eight blocks was randomized within each run, with no break between blocks. Four dummy images were collected after the last fixation cross. As a result, each run contained 84 images and lasted for 168 seconds (8 blocks x 20 seconds + 2 seconds x 4 dummy images).

**Fig. 3:**
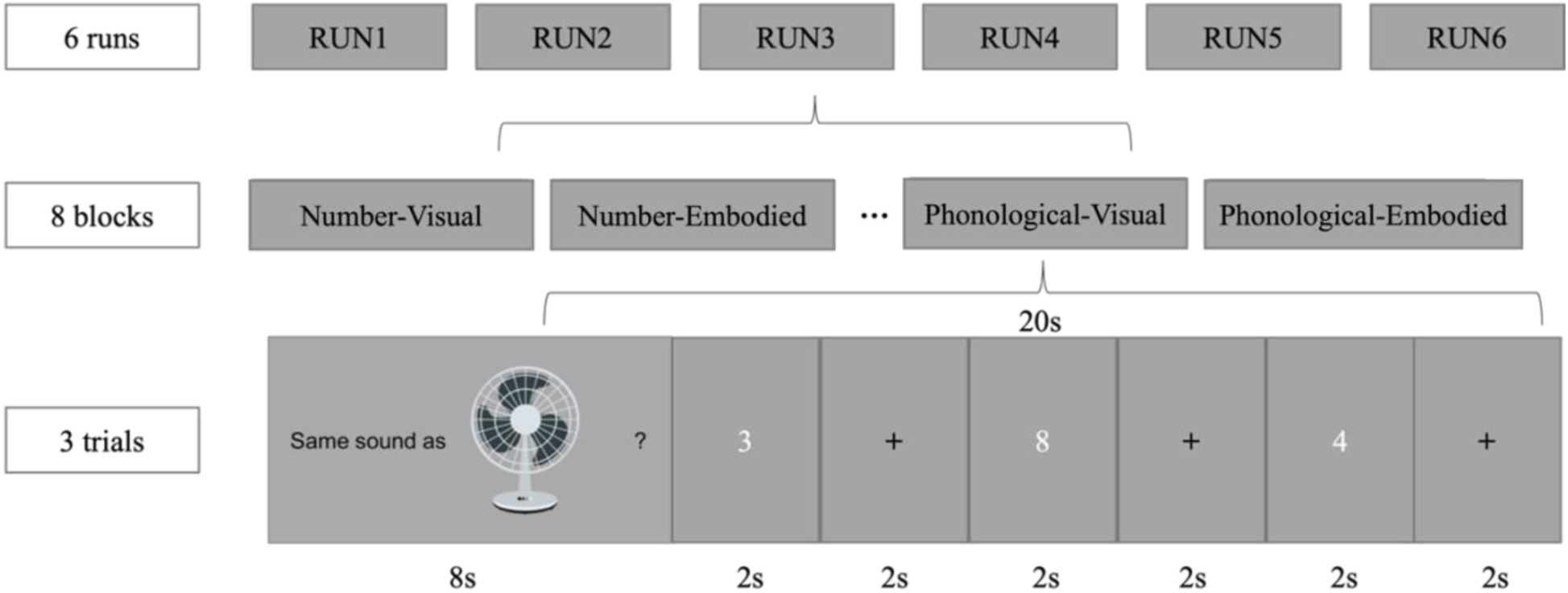
Experimental design of the fMRI task. Task included six runs, eight blocks in each run, with two blocks of each of the four conditions per run. Each block started with a prompt question, followed by three trials of the same condition (here: Phonological-Visual).

### 2.3 fMRI procedure

Children first completed a one-hour training session practicing the localizer task until they answered at least 75% of trials correctly and familiarizing themselves with the scanning environment and practicing to remain still in the scanner.

Both child and adult participants went through the same data acquisition procedures. They first completed three runs of resting state data each lasting for five minutes. These data were not of interest to the present study and are hence not reported here. Next, participants’ structural images were acquired (around 10 minutes) while viewing a video of an underwater scene with fish or a cartoon (for children), followed by the six functional runs described above. The entire experiment lasted around one hour; breaks were provided between runs and different data acquisition steps.

### 2.4 Imaging data acquisition and preprocessing

#### 2.4.1 Data acquisition

Data acquisition extended across two scanners. A 3T Siemens Verio MR scanner was used to acquire data from all adults and 26 children, while a 3T Siemens Prisma scanner was used for the remaining 62 children. The same imaging protocol was applied across both scanners. Specifically, both scanners were equipped with a 32-channel head coil and foam padding for head stabilization. Functional images were obtained using a T2* weighted echo-planar imaging (EPI) pulse sequence (TR = 2000ms, TE = 30ms, flip angle = 79 degrees, voxel size = 2 x 2 x 2mm, matrix = 106 x 106, 69 axial slices). T1-weighted anatomical scans were obtained in the middle of the imaging acquisition (TR = 2,330ms, TE = 1.97ms, flip angle = 9 degrees, voxel size = 1 x 1 x 1mm, matrix = 256 x 256, 176 sagittal slices).

#### 2.4.2 Imaging data preprocessing

We used the Analysis of Functional NeuroImages (AFNI) software package (Cox, 1996) to preprocess the imaging data. All functional images were slice-time corrected and were volume-registered to a mean functional volume. Functional images were normalized to standard Montreal Neurological Institute (MNI) space (Fonov et al., 2009) and smoothed with a Gaussian kernel of 6 mm full-width-half-maximum (FWHM). Motion was measured in six dimensions (rotation and shifts in x, y, z dimensions). Imaging data processing and analyses were identical for children and adults.

### 2.5 Behavioral assessments

Participants’ standardized math and reading abilities were tested in a separate behavioral session using the nationally normed Woodcock-Johnson III Tests of Achievement (Woodcock et al., 2001). The tests included five subtests: Calculation, Math Fluency, Applied Problems, Letter-Word Identification, and Word Attack. Participants also completed behavioral assessments measuring the integration between different number codes.

The Calculation subtest measures participants’ mathematical computation skills. It requires participants to solve problems that involve arithmetic, geometry, trigonometry, geometry, logarithms, and calculus. However, for children, the majority of the questions are arithmetic problems suitable for their ability levels. The Math Fluency subtest is a timed test that measures participants’ ability to solve as many simple addition, subtraction, and multiplication problems as possible within three minutes. The Applied Problems subtest requires participants to analyze and solve math word problems. During this subtest, the experimenter first verbally reads word problems to the participants, then participants select relevant information, recognize procedures, perform necessary calculations to get an answer, and then give answers verbally. A standard score across these three math subtests was calculated to represent participants’ overall math ability, which we used for the brain-behavioral correlational analyses.

The Letter-Word Identification subtest requires participants to identify letters and verbally read individual words correctly. The Word Attack subtest requires participants to first identify and produce sounds for single letters, and then verbally read aloud phonically regular nonwords. It is designed to measure participants’ ability to apply phonic and structural analysis skills to reading and decoding unfamiliar words. A standard score across Letter-Word Identification and Word Attack was calculated to represent participants’ overall basic reading skills.

To test participants’ integration between quantities and Arabic numerals, they completed a number comparison task featuring dot arrays and Arabic numerals in four different conditions (dot-dot, numeral-numeral, dot-numeral and numeral-dot). Numbers were sequentially presented and participants had to indicate whether the first or the second represented the larger quantity regardless of stimulus format. As in previous work (Ren et al., 2022), we calculated the response time difference between mixed trials (numeral-dot) and pure conditions (dot-dot, numeral-numeral) as a measure of symbolic estrangement.

### 2.6 Imaging data analysis

#### 2.6.1 Univariate analyses

We conducted a GLM to identify brain regions involved in number processing. First, for each participant, trials with no response, response times below 300 ms, and invalid button presses were interpolated with the mean response time for the respective condition. Subsequent univariate analyses involved applying GLMs to assess task contrasts (number > phonological task) within each stimulus type (Arabic numeral, hand) for each participant. The GLM involved convolving task onset times with a hemodynamic response function (HRF) using the dmBLOCK function in AFNI, with the HRF’s duration being modulated by the response time. Motion parameters were included in the model as nuisance regressors. Following the subject-level analysis, the estimated beta values from the GLMs were further submitted to a group-level analysis using the 3dttest++ function in AFNI. Voxels passing a significance threshold of *p* < .001 and clusters passing an alpha threshold (< .05) were selected from the group-level results. The conjunctions of the significant clusters for the task contrasts for the two stimulus types were then calculated, representing brain areas more responsive during number processing compared to phonological processing with either type of stimulus. These clusters were saved as masks for children and adults, respectively, and used for subsequent analyses.

#### 2.6.2 Brain-behavioral correlation analysis

In this study, we were interested in whether the activation of number processing brain regions correlated with participants’ math abilities. All 104 adult participants were included in the correlational analyses. One child was excluded due to missing data on math assessments, leaving 87 children in the correlational analyses. In addition, data from participants that were more than three standard deviations from the mean for each measurement (i.e., brain activation or math skills) were excluded (see Results section for details).

For both children and adults, we first extracted each significant cluster from the univariate analyses as a mask, and then applied the mask to each participant’s GLM results to calculate the mean beta values using the *3dROIstats* function in AFNI. The mean beta values represent the overall mean brain activation in each brain region. For children, we also applied the brain masks derived from adults to extract the mean brain activation within adult-matched regions. This approach allowed us to explore the relevance of these adult brain regions to the math abilities of children. We first calculated the Pearson correlation between the overall mean activation across all brain regions for each age group and participants’ math scores. Where significant correlations were found, we then proceeded to analyze each brain region individually, correlating the mean brain activation in each brain area with math scores. For regions that showed significant correlations with math abilities, we further explored their specificity by correlating the mean brain activation with reading skills.

Finally, we examined if brain activation in number processing brain regions also correlates with a behavioral index of symbolic estrangement derived from the number comparison task in adult participants. Ten adults were excluded due to missing comparison task data, which left 94 adults in this analysis. Since we did not find correlations between symbolic estrangement indices and math performance in children, we conducted this additional analysis only with adults’ data.

#### 2.6.3 Relations between activation of IPS and math abilities

Given the importance of the IPS in the literature and since the brain regions identified in our number localizer task (i.e., sensorimotor or inferior parietal areas) did not completely align with the IPS, we specifically explored the correlations between activation in the left and right IPS and adults’ and children’s math abilities. The exact locations of the IPS were derived from a study by Duricy et al.(2025), which used meta-analytic data from the Neurosynth database (Yarkoni et al., 2011). Specifically, the right and left IPS regions were generated based on 1) the Neurosynth v5-topics-100 meta-analysis Topic 18, featuring studies that included terms such as “magnitude,” “estimation,” and “number”; and 2) targeted searches for the terms “arithmetic” and “calculation.” Then, the authors created the IPS regions by identifying the largest parietal clusters in the left and right hemispheres across the three Neurosynth statistical maps, selecting voxels that overlapped in all three maps (Fig. 4). Thus, the IPS regions were identified with substantial evidence of involvement in number processing and math. Similar to our other correlational analysis, we applied the left and right IPS regions separately to adult and child data, extracted their mean brain activations (i.e., the mean beta values from the GLM with the number greater than phonological task contrast), and calculated correlations with the overall math abilities in both children and adults.

**Fig. 4:**
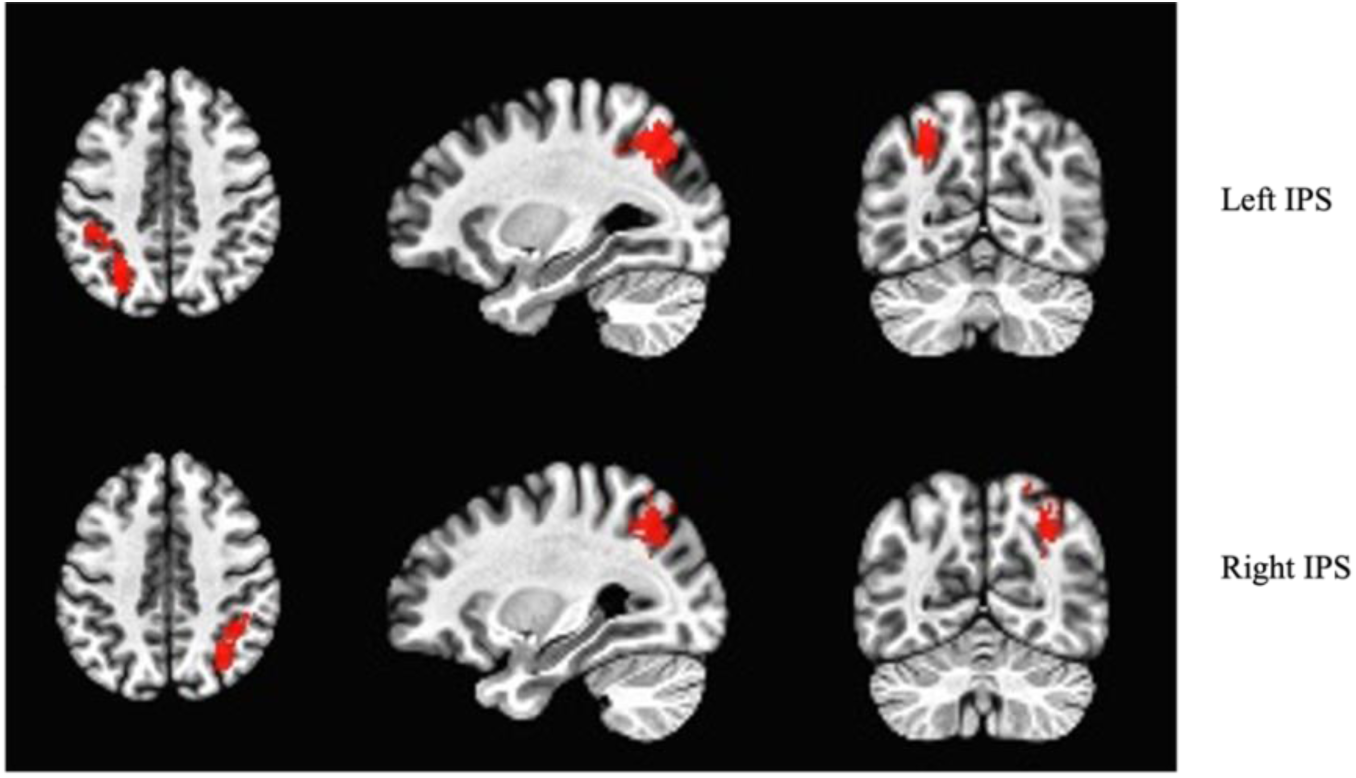
IPS regions from the study by Duricy et al.(2025).

## 3. Results

### 3.1 Adults: Univariate analysis of imaging data

For adults, the univariate analysis revealed six clusters that were more active during the number task compared to the phonological task. The six clusters encompass tissue that localizes to: 1) the right occipital/temporal/inferior parietal cortices, 2) the left occipital/temporal cortices, 3) the left somatosensory/motor cortices, 4) the right somatosensory cortex/supramarginal gyrus, 5) the bilateral supplementary motor/motor cortices, and 6) the right insular cortex (see Fig. 5 and Table 1 for details).

**Fig. 5:**
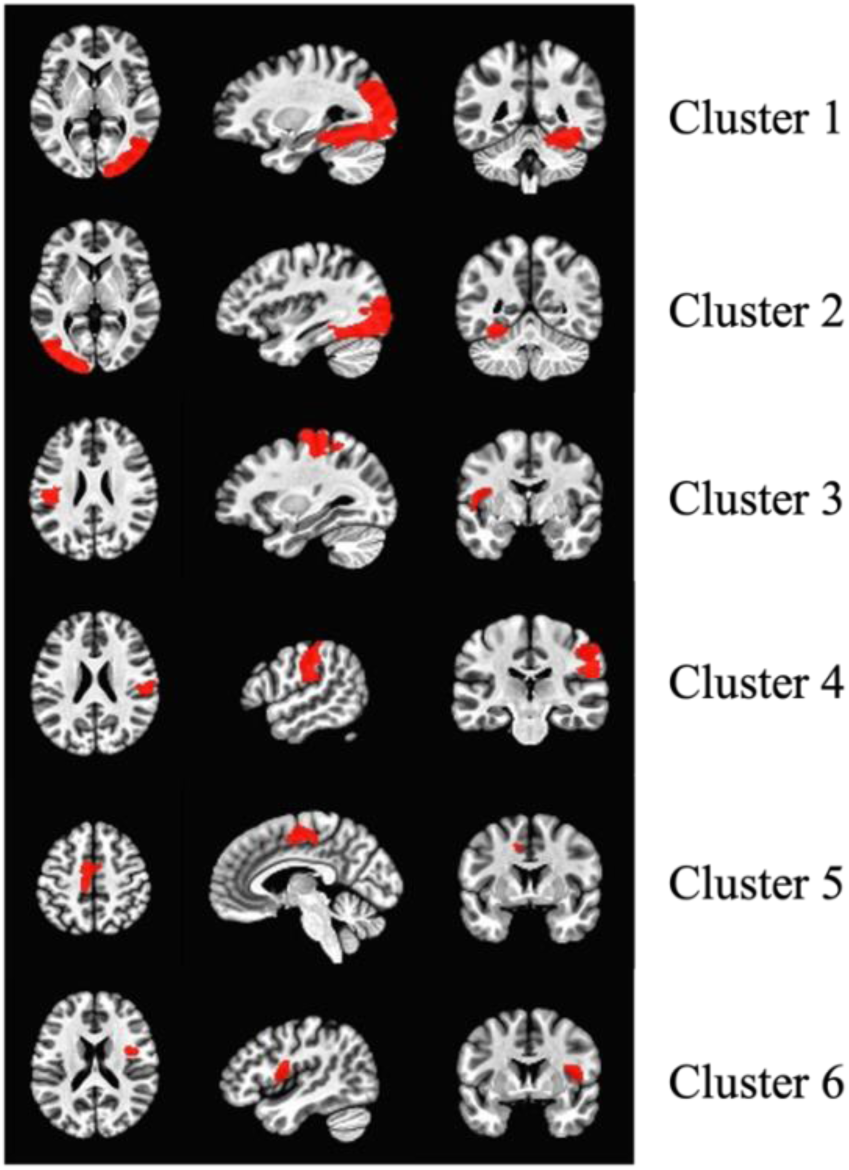
Univariate analysis results for adults. Brain areas involved in number processing in MNI space with reference to Broadman’s areas (BA). Only BAs with more than 100 significantly active voxels are noted here. Cluster 1: Right occipital (BA17/BA18/BA19)/temporal (BA37/BA20)/inferior parietal (BA39) cortices. Cluster 2: Left occipital (BA17/BA18/BA19)/temporal (BA37/BA20) cortices. Cluster 3: Left somatosensory (BA1/BA2/BA3)/motor (BA4) cortices. Cluster 4: Right somatosensory cortex (BA2)/supramarginal gyrus (BA40). Cluster 5: Bilateral supplementary motor (BA6)/left motor (BA4) cortices. Cluster 6: Right insular cortex.

**Table 1:** Univariate analysis results for adults. L indicates left hemisphere, and R indicates right hemisphere. The coordinates (x, y, z) represent the center of each cluster in MNI space. Cluster 1: Right occipital (BA17/BA18/BA19)/temporal (BA37/BA20)/inferior parietal (BA39) cortices. Cluster 2: Left occipital (BA17/BA18/BA19)/temporal (BA37/BA20) cortices. Cluster 3: Left somatosensory (BA1/BA2/BA3)/motor (BA4) cortices. Cluster 4: Right somatosensory cortex (BA2)/supramarginal gyrus (BA40). Cluster 5: Bilateral supplementary motor (BA6)/left motor (BA4) cortices. Cluster 6: Right insular cortex.

| Clusters | Hemisphere | # of voxels | x | y | z |
| --- | --- | --- | --- | --- | --- |
| Cluster 1 | R | 7392 | 32 | -72 | 0 |
| Cluster 2 | L | 5740 | -28 | -80 | -4 |
| Cluster 3 | L | 3065 | -40 | -24 | 44 |
| Cluster 4 | R | 1296 | 52 | -24 | 36 |
| Cluster 5 | L/R | 1018 | -4 | -12 | 52 |
| Cluster 6 | R | 319 | 40 | 0 | 16 |

### 3.2 Children: Univariate analysis of imaging data

For children, the univariate analysis revealed three clusters with greater activation during the number task compared to the phonological task. The three clusters encompass tissue that localizes to: 1) the left occipital cortex, 2) the right supramarginal gyrus, and 3) the right inferior parietal cortex (see Fig. 6 and Table 2 for details).

**Fig. 6:**
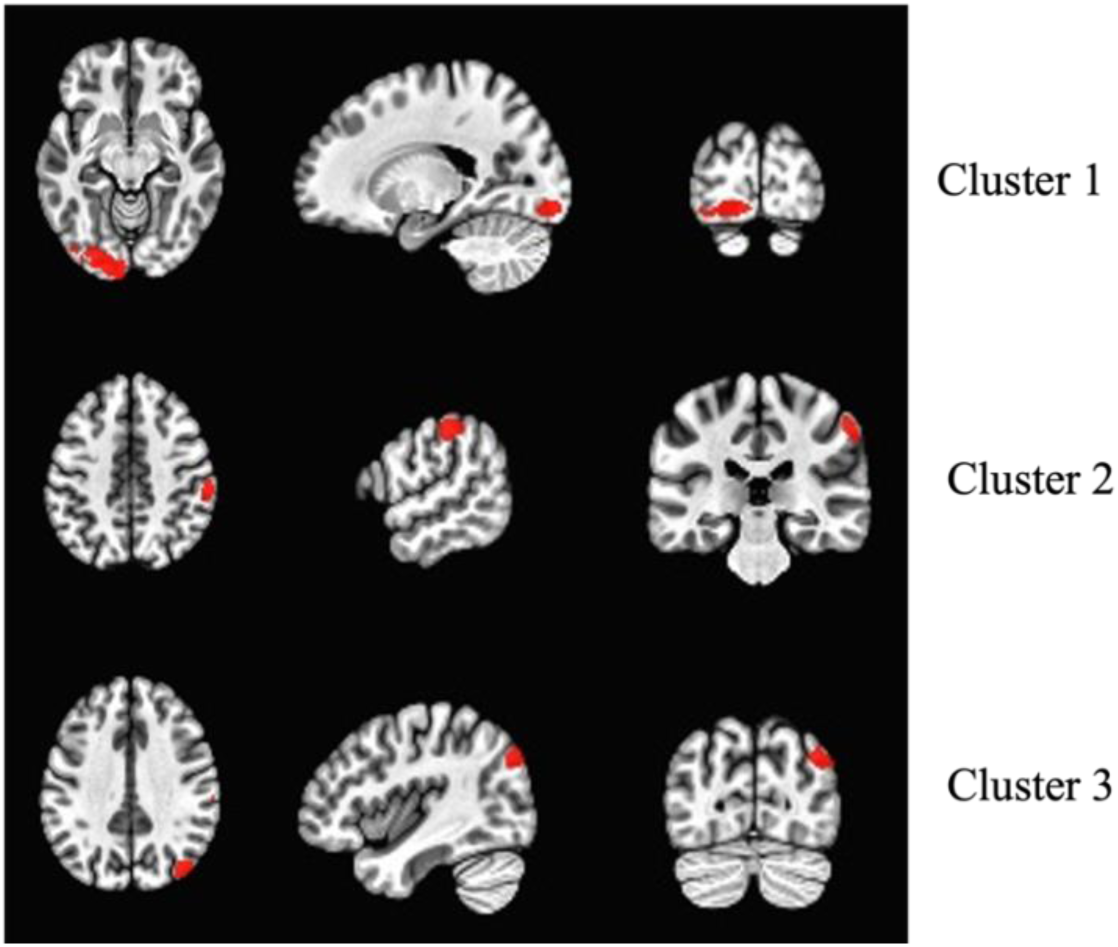
Univariate analysis results for children. Brain areas involved in number processing in MNI space with reference to Broadman’s areas (BA). Only BAs with more than 100 voxels are noted here. Cluster 1: Left occipital cortex (BA18/19). Cluster 2: Right supramarginal gyrus (BA40). Cluster 3: Right inferior parietal cortex (BA39).

**Table 2:**
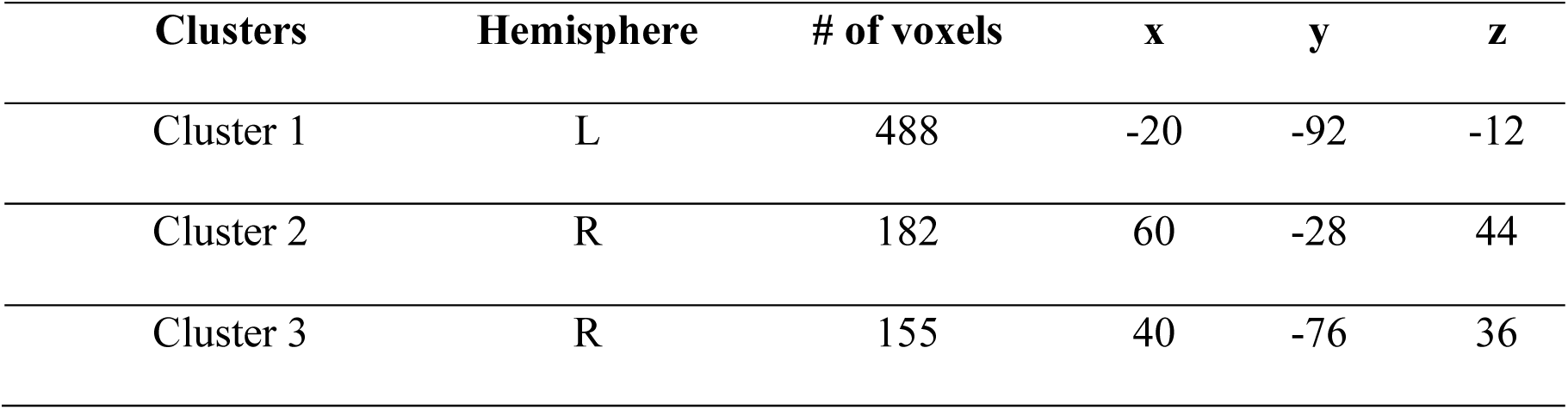
Univariate analysis results for children. L indicates left hemisphere, and R indicates right hemisphere. The coordinates (x, y, z) represent the center of each cluster in MNI space. Cluster 1: Left occipital cortex (BA18/19). Cluster 2: Right supramarginal gyrus (BA40). Cluster 3: Right inferior parietal cortex (BA39).

### 3.3 Brain-Behavioral correlational analyses

#### 3.3.1 Adults

Four adults were excluded from the brain-behavior correlations due to brain activation beyond three standard deviations from the group mean, and one adult was excluded due to math ability exceeding three standard deviations from the group mean. Thus, a total of 99 subjects remained in the correlational analyses.

For the correlation analyses, we found a significant negative correlation between adults’ math skills and the mean brain activation during the number task relative to the phonological task across the six brain areas identified in the univariate analysis above, *r* (97) = -.24, *p* = .02. Examining each of the six areas separately, we found significant negative correlations between adults’ math abilities and left somatosensory/motor cortices (*r* (97) = -.27, *p* = .007), right somatosensory cortex/supramarginal gyrus (*r* (97) = -.22, *p* = .03), and bilateral supplementary motor/left motor cortices (*r* (97) = -.21, *p* = .03; see Fig. 7). The correlation between adults’ math abilities and right insular cortex activation was marginally significant, *r* (97) = -.20, *p* = .05. However, no significant correlations were observed between math abilities and right occipital/temporal/inferior parietal cortices (*r* (97) = -.14, *p* = .17) or left occipital/temporal cortices (*r* (97) = -.13, *p* = .21).

**Fig. 7.**
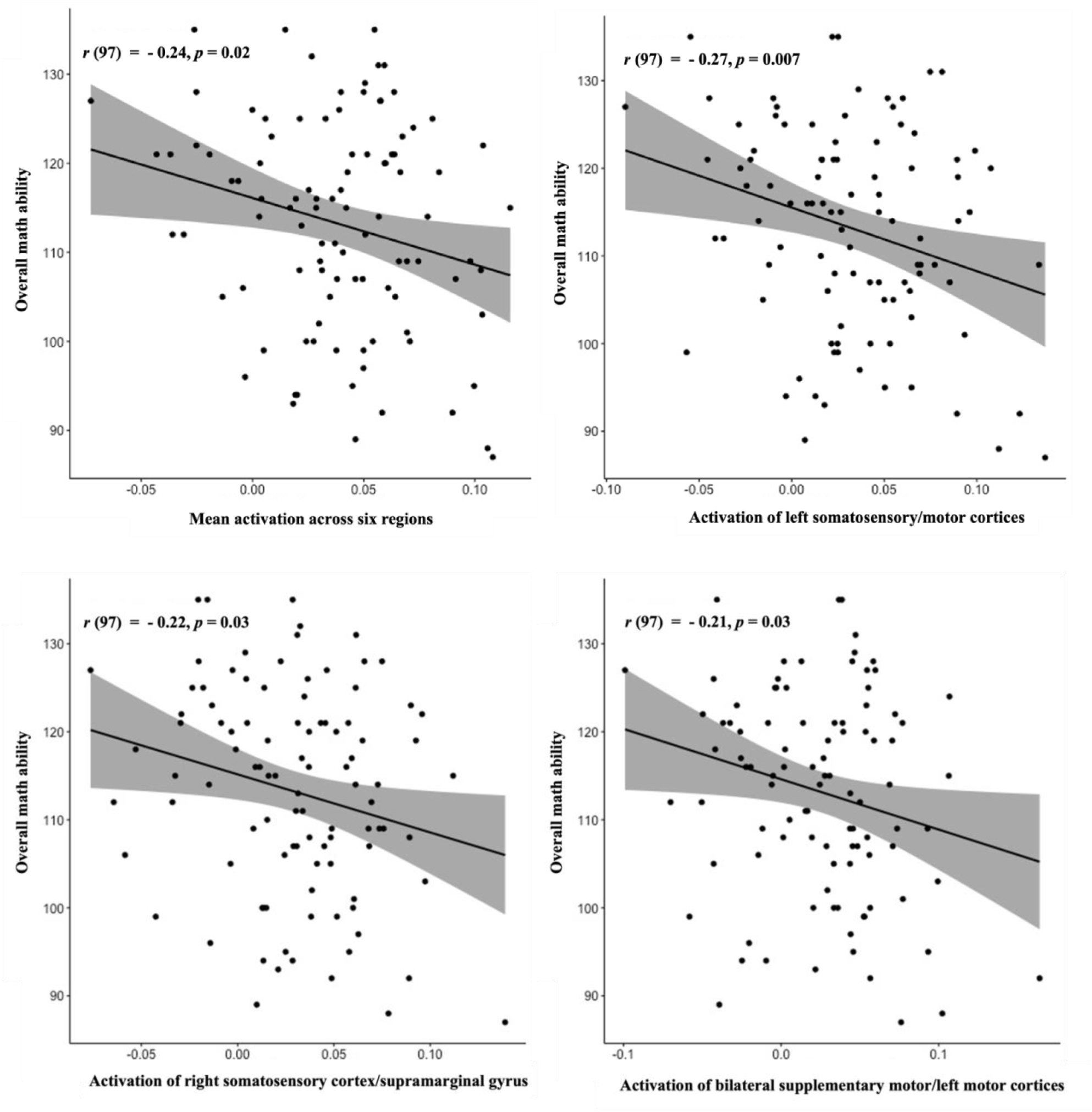
Scatterplots depicting the association between mean brain activation differences for the number task relative to the phonological task across six number processing regions in adults (Table 1) (top left) and adults’ math ability score on the math subtests of the Woodcock-Johnson III Tests of Achievement. The three remaining panels depict the association for specific brain regions: left somatosensory/motor cortices (top right), right somatosensory cortex/supramarginal gyrus (bottom left), and bilateral supplementary motor/left motor cortices (bottom right). Solid black lines represent regression line, gray areas represent 95% confidence interval.

For the three brain areas that showed significant correlations with math abilities, we also tested the specificity of these correlations by examining their relations with reading skills (see Appendix figure 1 for reading score distribution). One additional subject was excluded due to their reading score exceeding three standard deviations from the mean. We did not find significant correlations with reading ability in the left somatosensory/motor cortices, right somatosensory cortex/supramarginal gyrus, and bilateral supplementary motor/left motor cortices (all *ps* > .25).

Finally, we examined the correlation between mean brain activation during the number task relative to the phonological task across all six brain areas and adults’ symbolic estrangement index. One adult was excluded due to mean brain activation being beyond three standard deviations from the group mean, and another was excluded due to a symbolic estrangement index exceeding three standard deviations from the mean, leaving 92 adults in the analysis. We did not find a significant correlation between the mean brain activation and symbolic estrangement, *r* (90) = .02, *p* = .83.

#### 3.3.2 Children

Two children were excluded due to mean brain activation exceeding three standard deviations from the group mean, and another two were excluded due to math ability being beyond three standard deviations from the group mean. Thus, data from a total of 83 children remained in the correlational analyses. We did not find significant correlations between mean activation during the number task relative to the phonological task across three brain areas in children and their math ability (*r* (81) = .06, *p* = .57), or between mean brain activation across the six adult-defined regions and children’s math ability (*r* (81) = .10, *p* = .35). Thus, no further analyses were warranted.

### 3.4 Brain activation of IPS and math abilities

To examine the role of the IPS for adults’ and children’s math abilities, we correlated the brain activation in left and right IPS during the number task relative to the phonological task and math abilities in both children and adults. For adults, one participant was excluded due to brain activation being beyond three standard deviations from the group mean, which resulted in a final analytic sample of n = 103. We found a significant negative correlation between brain activation in the left IPS and adults’ math abilities, *r* (101) = -.26, *p* = .008 (Fig. 8). However, there was no significant correlation between activation in the right IPS and math abilities in adults, *r* (101) =-.11, *p* = .25.

**Fig. 8:**
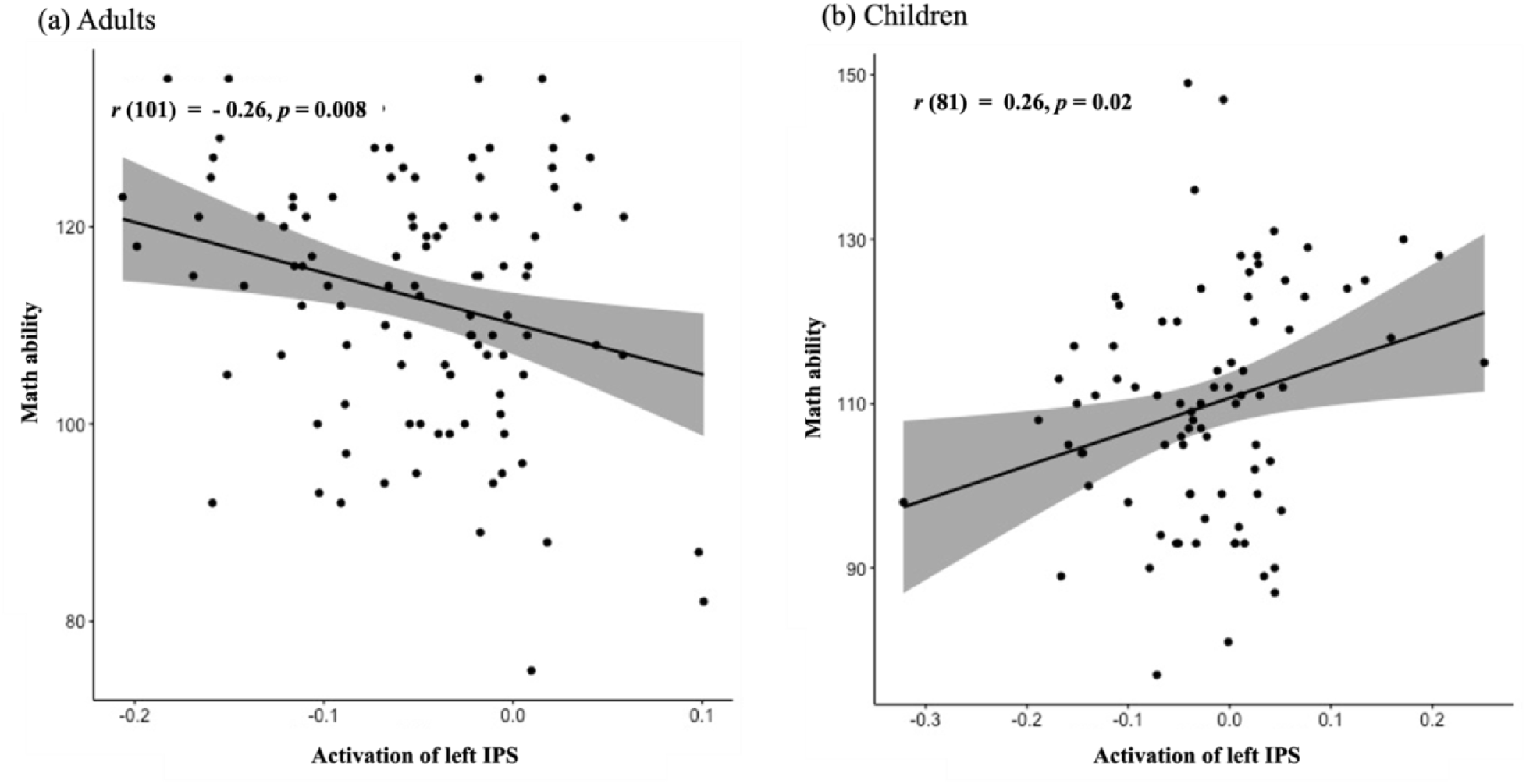
Scatterplots depicting the association between mean brain activation during the number task relative to the phonological task in the left IPS and adults’ (left panel) and children’s (right panel) math ability score on the math subtests of the Woodcock-Johnson III Tests of Achievement. Solid black lines represent regression line, gray areas represent 95% confidence interval.

For children, two participants were excluded due to brain activation exceeding three standard deviations from the group mean, and another two due to math abilities exceeding three standard deviations from the mean. Thus, a total of 83 children remained in the analyses. There was a significant positive correlation between brain activation in the left IPS and math abilities in children, *r* (81) = .26, *p* = .02 (Fig. 8). Similar to adults, we did not find a significant correlation between the brain activation in the right IPS and math abilities in children, *r* (81) = .17, *p* = .13.

To test the specificity of these results, we examined the correlations between activation of the left IPS and reading skills in adults and children. For adults, one additional participant was excluded due to reading skill exceeding three standard deviations from the group mean, leaving 102 participants in the analyses. We did not find a significant correlation between activation of the left IPS and reading skills in adults, *r* (100) = -.05, *p* = .66. Similarly in children, we also did not find a significant correlation between brain activation of the left IPS and reading skills, *r* (81) = .18, *p* = .10.

## 4. Discussion

In our study, we used a novel number localizer task featuring numeral (symbolic) and hand (embodied) number codes to characterize the neural underpinnings of number processing in both children and adults. For adults, we identified a broader set of brain areas including occipital, temporal, parietal, and the insular cortices that were more active during the number compared to the phonological task. For children, brain areas more active during number compared to phonological processing were primarily in the occipital cortex, supramarginal gyrus, and inferior parietal cortex. These findings suggest that adults engage a broader set of brain regions compared to children, aligning with previous literature that indicates individuals with higher math proficiency tend to activate a more extensive neural network (Zamarian et al., 2009). Engaging a wider array of neural resources may enhance adults’ cognitive processing and neural efficiency (O’Boyle et al., 2005).

Interestingly, *reduced* activation in somatosensory/motor areas was associated with *better* math competence in adults. In light of the theory of embodied numerical cognition, which posits that the understanding of numbers is fundamentally linked to sensorimotor experiences, such as finger counting or spatial interactions (Barsalou, 2008; Sixtus et al., 2023), our findings indicate that less reliance on these processes is associated with higher math skills. However, this does not mean that the embodied foundation completely disappears in adulthood. Previous studies using a range of neuroimaging techniques, including fNIRS, has identified representations of numerical information in sensorimotor areas (Artemenko et al., 2022). Research consistently shows that childhood finger-counting strategies continue to influence adults’ mental representation of numbers (Di Luca et al., 2006; Di Luca & Pesenti, 2008), and that these finger–numeral representations may also influence simple arithmetic operations (Badets et al., 2010). This bodily experience has long-lasting effects in shaping the structure of adult number processing (Domahs et al., 2010). Our findings further suggest that while the embodied foundation persists, relying less on this processing is associated with higher math abilities in adults. This pattern of reduced activation with increased skill is a well-documented hallmark of learning-related neural efficiency, not unique to numerical cognition. For example, in motor sequence learning, activation levels consistently decrease in related cortical regions as performance becomes more automatic, reflecting more efficient processing (Reithler et al., 2010). Similarly, motor areas adaptively reduce activity for learned sequences, marking a transition from effortful control to automaticity (Poldrack et al., 2005). In line with neural efficiency theory, our findings suggest that, while embodied representations of number may be crucial at early developmental stages for grounding and supporting symbolic numerical knowledge, adults may not rely heavily on embodied number codes for higher processing efficiency. It is possible that with increased experiences with numbers, adults develop greater automaticity and access to symbolic representations, leading to a reduced reliance on embodied representations, i.e., reduced activation in motor areas, to facilitate a transition to more abstract number representations and efficient cognitive processes. In other words, while embodied representations may serve as a scaffold during early development, continued heavy reliance on them in later stages might hinder the transition to more abstract and automatic processing, which is essential for efficient math problem-solving. Thus, our findings do not argue that the embodied foundation has been completely abandoned in adulthood. Rather, they suggest that adult math proficiency is marked by a fundamental shift from effortful processing to efficient, automated processing. We did not find any associations between brain activation in sensorimotor areas and math abilities in fourth graders, but this may be due to the fact that children at this age and level of formal education in math do not consistently rely on embodied representations.

Regarding the role of the IPS, we found that activation in the left IPS correlates with math abilities in both children and adults, but we did not find such correlations in the right IPS. This pattern supports the developmental trajectory of the IPS, where the right IPS is more involved at earlier stages of numerical processing, observable in preverbal infants and children aged 3 to 6 years (Edwards et al., 2016; Kersey & Cantlon, 2017). The left IPS becomes increasingly dominant as individuals’ experience with math and numbers grows (Ansari, 2016; Emerson & Cantlon, 2015). Interestingly, we observed that less activation in the left IPS correlates with better math abilities in adults, whereas greater activation is linked with higher math abilities in children. This finding aligns with the neural efficiency hypothesis (O’Boyle et al., 2005), which suggests that experienced adults may rely less on basic quantity processing. In contrast, children, possibly due to their lack of experience with formal math, need to access magnitude information and rely on basic quantity processing to complete math tasks. This interpretation aligns with previous behavioral evidence indicating that adults with higher math skills are less likely to activate magnitude information, particularly during symbolic number processing, compared to those with poorer skills (Ren et al., 2022).

In conclusion, our findings support the notion that proficient math abilities in adults are characterized by reduced reliance on processing grounded in sensorimotor experience. This shift likely facilitates more efficient and abstract number processing and cognitive processes.

Moreover, our study highlights the important role of the left IPS and its differential involvement in math competence in adults and children. Adults may engage the IPS less as the need to access quantitative information becomes less critical compared to children (Ren et al., 2022). This pattern in adults likely supports a more abstract number processing and more efficient neural processing (Ren & Libertus, 2023).

In summary, our research sheds light on the neural underpinnings of number processing and math competence in both children and adults. Importantly, our findings raise questions about the emphasis on embodied number representations in math learning experiences. While these representations may serve as a scaffold in the early stages, as individuals develop greater automaticity in number processing, they may rely less on embodied representations, facilitating the use of more abstract number representations and cognitive processes for efficient math problem-solving. It is important to carefully consider this progression when developing math learning strategies, particularly for individuals who face challenges in learning math, to better address diverse learning needs. Moreover, future research should delve deeper into investigating *when* and *how* this progression occurs. Longitudinal studies could serve as an initial step in these endeavors, providing us with deeper insights.

## Acknowledgement

This work was supported by the National Science Foundation (grant number: 1734735). We would like to thank Taylor Casteel and Corrine Durisko for their help. We also would like to thank Erin Duricy and colleagues for kindly providing the IPS regions from their study.

## Appendix

**Fig S1:**
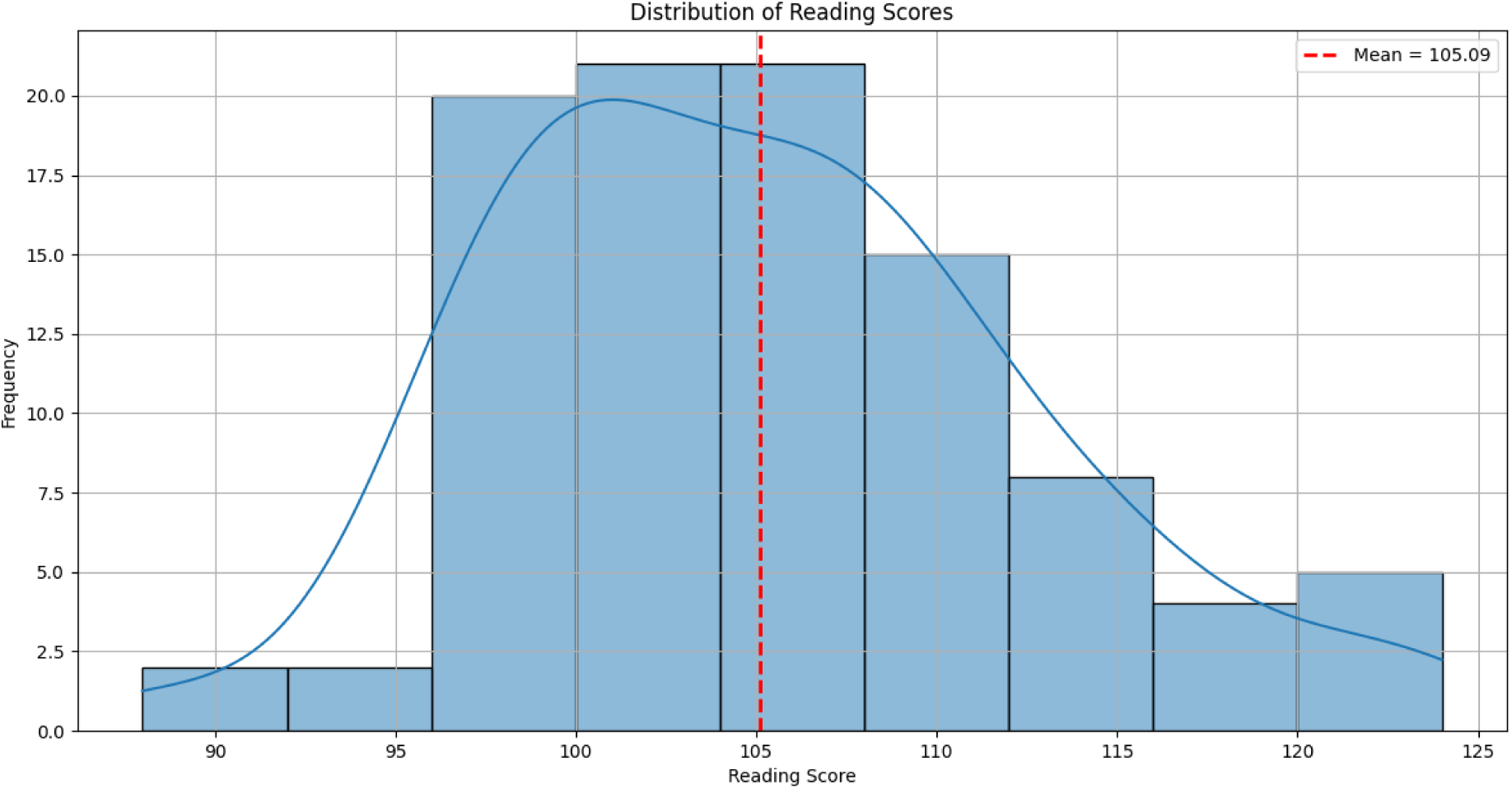
Distribution of reading scores, a standard score from the Letter-Word Identification and Word Attack subtests representing participants’ overall reading skills.

## Notes

### Competing Interest Statement

The authors have declared no competing interest.

